# TPR is required for cytoplasmic chromatin fragment formation during senescence

**DOI:** 10.1101/2024.04.18.590085

**Authors:** Bethany M. Bartlett, Yatendra Kumar, Shelagh Boyle, Tamoghna Chowdhury, Andrea Quintanilla, Charlene Boumendil, Juan Carlos Acosta, Wendy A. Bickmore

## Abstract

During oncogene-induced senescence there are striking changes in the structure of the nucleus and the organisation of heterochromatin. This is accompanied by activation of a pro-inflammatory gene expression programme – the senescence associated secretory phenotype (SASP) – driven by transcription factors such as NF-κB. Here we show that TPR, a protein of the nuclear pore complex basket, is required for the very early activation of NF-κB signalling during the stress-response phase of oncogene-induced senescence. This is prior to activation of the SASP and occurs without affecting NF-κB nuclear import. We show that TPR is required for the activation of TBK1 signalling at these early stages of senescence and we link this to the formation of heterochromatin-enriched cytoplasmic chromatin fragments thought to bleb off from the nuclear periphery. These cytoplasmic chromatin fragments appear to lack nuclear pore components. Our data suggest that TPR at the nuclear pore is involved in the loss of structural integrity of the nuclear periphery during senescence. We propose that this acts as a trigger for activation of cytoplasmic nucleic acid sensing, NF-κB signalling, and activation of the SASP, during senescence.

## Introduction

DNA damage, such as short telomeres (replicative senescence) or oncogene signalling, can trigger senescence, an irreversible cell cycle arrest programme. During oncogene-induced senescence (OIS) chromatin organisation is dramatically disrupted. Pre-existing heterochromatin moves away from the nuclear periphery (Chandra et al., 2012), forming internal senescence-associated heterochromatic foci (SAHF) (Narita et al., 2003).

Senescent cells also activate a gene expression programme that leads to the secretion of a cocktail of inflammatory cytokines, chemokines, and growth factors - known as the senescence-associated secretory phenotype (SASP) (Coppé et al., 2010; Acosta et al., 2013). The SASP can contribute to tumour suppression by enhancing immune cell recruitment (Kale et al., 2020; Xue et al., 2007), but it can also promote tumour growth (Kuilman et al., 2008), and immunosuppression (Ruhland et al., 2016). Activation of SASP-related genes is primarily driven by the transcription factors (TFs) NF-κB (subunit p65) and C/EBPβ (Chien et al., 2011; Kuilman et al., 2008) and is accompanied by substantial changes in the landscape of active enhancers (Martínez-Zamudio et al., 2020; Tasdemir et al., 2016).

As well as transitioning to the nuclear interior to form SAHF, heterochromatin blebs off from the nuclear membrane during OIS, forming cytoplasmic chromatin fragments (CCFs) (Ivanov et al., 2013). CCFs are enriched for the heterochromatin-associated histone modifications H3K9me3 and H3K27me3 (Dou et al., 2017; Ivanov et al., 2013). CCFs are also positive for the DNA damage marker γ-H2AX, suggesting that DNA damage may play a role in CCF formation (Ivanov et al., 2013). In the cytoplasm, CCFs are sensed by the cGAS-STING pathway, which leads to activation of the SASP via NF-κB signalling (Dou et al., 2017; Glück et al., 2017; Yang et al., 2017).

The density of nuclear pores increases during OIS and proteins of the nuclear pore complex (NPC) play a role in both the formation of SAHF and activation of the SASP (Boumendil et al., 2019). TPR is a 267-kDa protein present in the nuclear basket (Figure 1A), which anchors to the NPC through its interaction with NUP153 (Hase and Cordes, 2003). TPR consists of four coiled-coil domains at its N-terminus, and an acidic C-terminal domain (CTD), which is phosphorylated by ERK2 (Vomastek et al., 2008). TPR is necessary for heterochromatin exclusion at nuclear pores (Krull et al., 2010) and we have previously shown that TPR anchored at the nuclear pore is necessary for both the formation and maintenance of SAHF, as well as for activation of the SASP, during OIS without affecting cell cycle exit (Boumendil et al., 2019). Here we investigate the requirement of TPR for SASP activation during OIS as well as during the early replicative stress that occurs in response to oncogenic RAS induction. Our results suggest a key role for TPR in the activation of innate immune signalling linked to the formation of cytoplasmic chromatin fragments.

**Figure 1.**
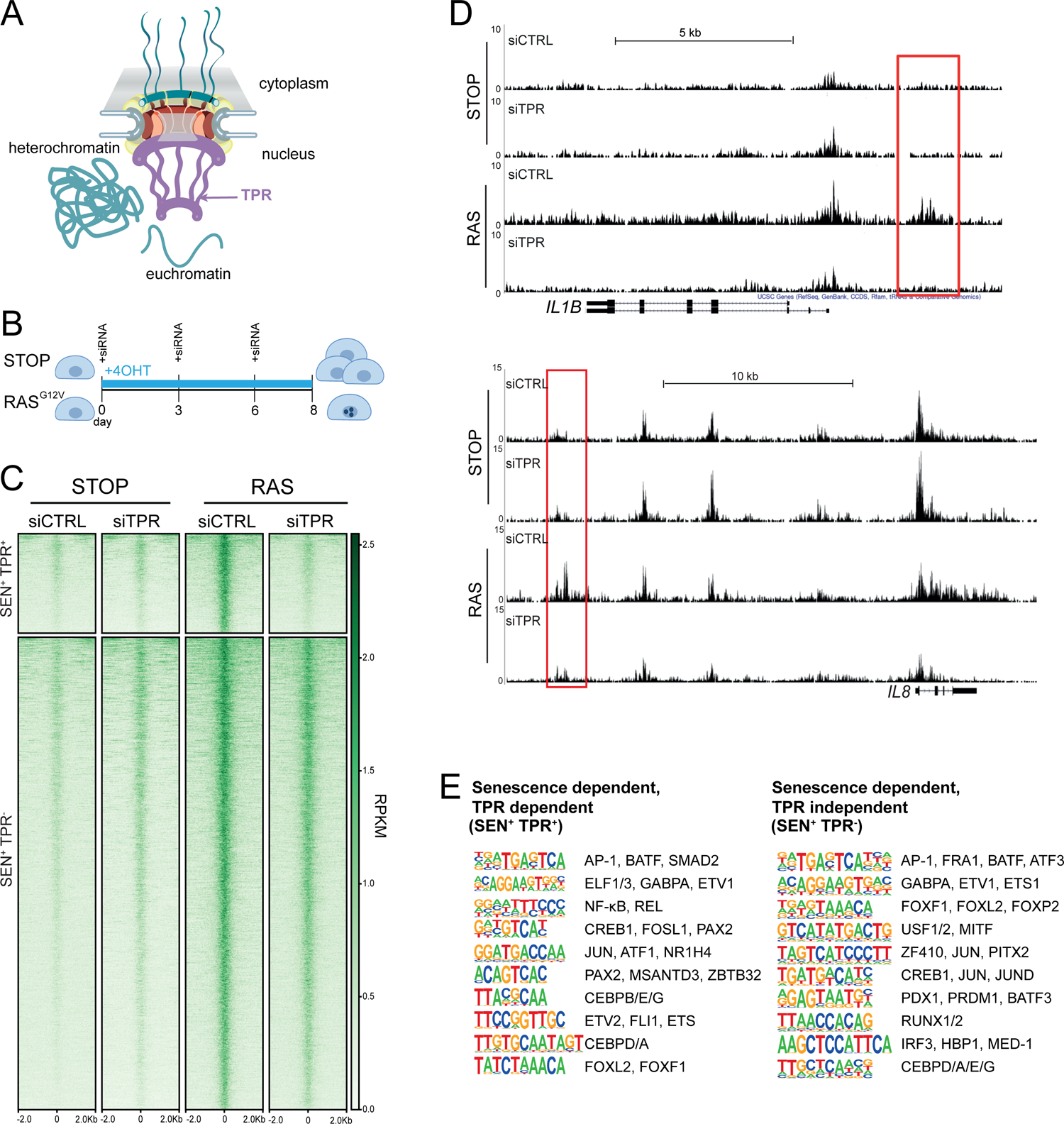
Senescence specific enhancers dependent on TPR are near SASP genes and are enriched in binding sites for $ASP-related transcription factors. A) Model of the nuclear pore showing the location of TPR in the nuclear basket and heterochromatin exclusion at the pore. B) Schematic of experimental protocol for senescence induction in IMR90 cells. After 8 days of treatment with 4-hydroxytamoxifen (4-OHT), the control (STOP) line continues to proliferate while the RAS line becomes senescent due to induction of RAS^G12V^ expression. C) Heatmap showing ATAC-seq signal in control (STOP) and O1S (RAS) cells 8 days after treatment with 4-OHT and transfection with either control (CTRL) or TPR siRNAs. SEN^+^ indicates signal specific to senescent cells and TPR^+^ indicates dependence on TPR. Intensity scale represents Reads per Kilobase per Million mapped reads (RPKM). D) Track views of ATAC-seq data from STOP and RAS cells treated with CTRL or TPR siRNAs at *IL1B* (top) and */LB* (bottom) gene loci. E) HOMER motif analysis of the senescence and TPR dependent ATAC-seq peaks (SEN^+^ TPR^+^) and the peaks that are dependent on senescence but not TPR (SEN+ TPR-). The top ten motifs are shown for each category of peaks. For both analyses all motifs have a p-value < 10^-13^

## Results

### Enhancers dependent on TPR during senescence are enriched for binding sites of inflammatory transcription factors

To study the role of TPR in the activation of the SASP that follows OIS, we used IMR90 fibroblasts harbouring an estrogen-inducible (4-OHT) oncogenic RAS^G12V^ mutation (ER:RAS^G12V^) (Acosta et al., 2013). The enhancer landscape of IMR90 cells changes during OIS (Tasdemir et al., 2016) and there is evidence that some nucleoporins interact with enhancers and regulate the transcriptional activity of associated genes (Ibarra et al., 2016; Pascual-Garcia et al., 2017). Therefore, we investigated whether TPR influences the enhancers that regulate SASP gene activation. We used ATAC-seq to identify whether there are regions of accessible chromatin that are specific to senescent cells, and that are TPR-dependent – i.e. are lost after TPR depletion by siRNAs at day 8 (d8) of RAS-induced senescence (Figure 1B).

Of the 6826 peaks with a significant increase in accessibility in senescent (RAS siCTRL) compared to non-senescent control (STOP siCTRL) cells (senescent dependent (SEN^+^)), 1187 are also TPR dependent (SEN^+^TPR^+^) (Figure 1C, Figure S1A, Table S1). Many of these include putative enhancers located close to key SASP genes, such as *IL1B* and *IL8* (Figure 1D). When we plotted published ChIP-seq data for H3K27 acetylation (H3K27ac) during OIS in IMR90 ER:HRAS^G12V^ cells treated with 4-OHT for six days (Parry et al., 2018) against our ATAC-seq data, both SEN^+^TPR^+^ and SEN^+^TPR^-^ peak categories showed an increase in H3K27ac in senescent cells when compared with the non-senescent control (Figure S1B). This further suggests that the regions which become accessible upon senescence function as senescence-specific enhancers, regardless of their dependence on TPR.

**Figure S1.**
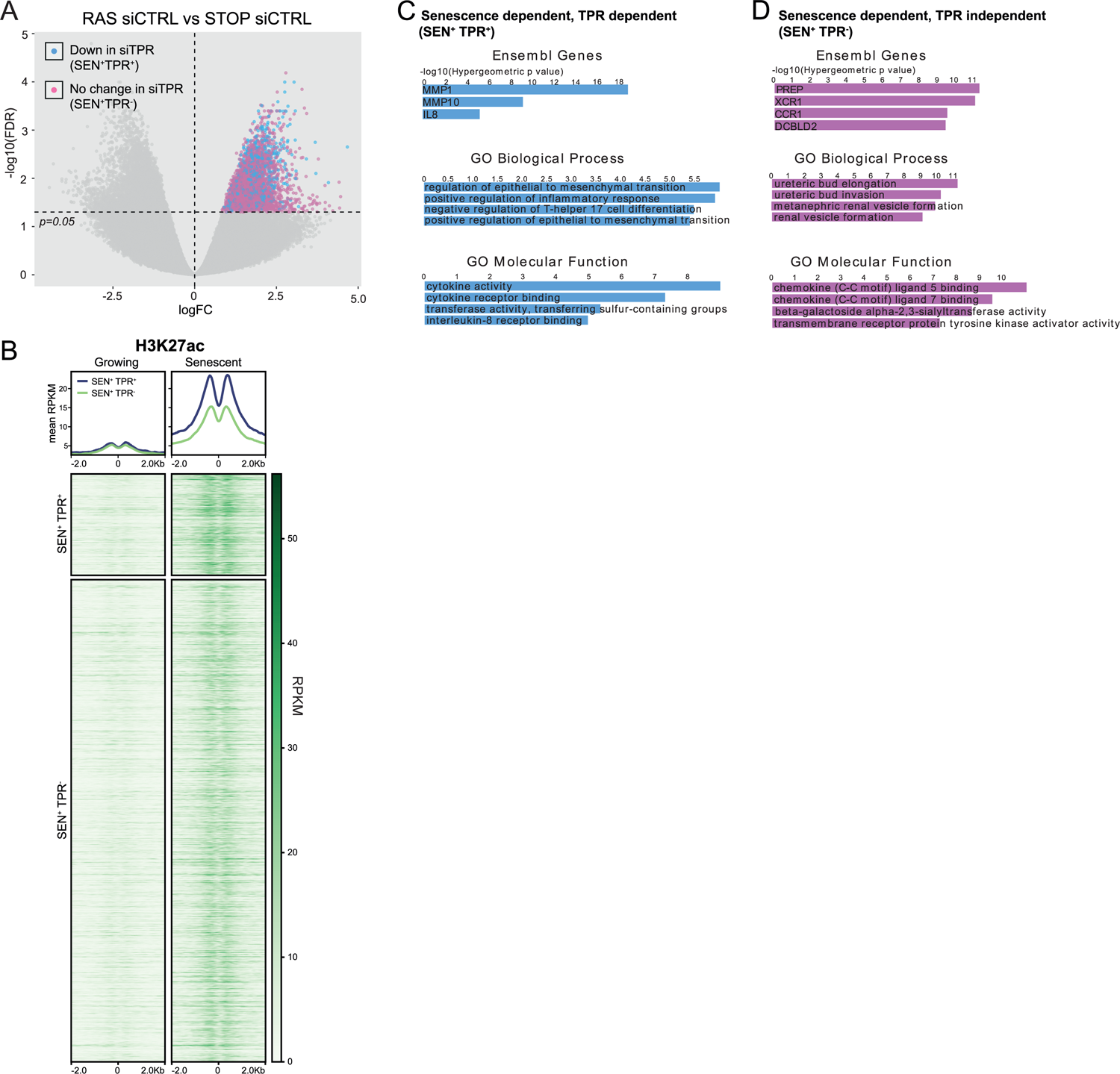
TPR dependent senescence specific enhancers are enriched in H3K27ac and associated with genes relevant to inflammation. Related to Figure 1. A) Volcano plot of differential accessibility analysis of d8 ATAC-seq peaks in RAS siCTRL vs STOP siCTRL. The horizontal dashed line indicates an adjusted p-value (FDR) of 0.05. Peaks with a significant increase in accessibility in senescent cells (SEN^+^) are coloured blue if their accessibility decreases on siTPR treatment (SEN^+^ TPR^+^), or pink if their accessibility does not change on siTPR treatment (SEN^+^ TPR). B) Heatmap of H3K27 acetylation (H3K27ac) ChIP-seq data from growing and senescent IMR90 cells (Parry et al., 2018) for the TPR dependent (TPR^+^) and TPR-independent (TPR) peaks defined in Figure 1C. C) Top ranking results of Gene Ontology analysis of genes close to SEN^+^ TPR^+^ peaks using the GREAT package (McLean et al., 2010). D) As in (C) but for genes close to SEN^+^ TPR-peaks.

Gene ontology (GO) analysis carried out using the Genomic Regions Enrichment of Annotations tool (GREAT) (McLean et al., 2010) showed that TPR-dependent peaks are significantly near to known SASP factor genes, and to genes enriched in Biological Process and Molecular Function categories such as ‘positive regulation of inflammatory response’, and genes involved in cytokine activity and cytokine receptor binding (Figure S1C). TPR independent senescent-dependent peaks showed proximity to chemokine receptor genes (*XCR1*, *CCR1*) (Figure S1D) whose expression allows cells to sense and respond to chemokines such as those secreted in the SASP (Coppé et al., 2010). However, the TPR independent peaks did not show proximity to any SASP factor genes, suggesting that senescence activated enhancers close to SASP genes (Tasdemir et al., 2016) may all be TPR dependent.

HOMER motif analysis (Heinz et al., 2010) revealed that d8 SEN^+^TPR^+^ enhancers are enriched for motifs for TFs, such as NF-κB and C/EBPβ, which are known to regulate the SASP (Acosta et al., 2008; Kuilman et al., 2008) (Figure 1E). TPR independent enhancers (SEN^+^TPR^-^) are not enriched for these motifs. This indicates that during OIS, TPR is involved in regulation of the NF-κB dependent proinflammatory SASP. Both categories of senescent dependent peaks are enriched in binding motifs for components of the AP-1 complex (Figure 1E). AP-1 acts as a pioneer TF, premarking prospective senescence enhancers (Martínez-Zamudio et al., 2020). This suggests that the initial shaping of the senescence enhancer landscape by AP-1 is unaffected by TPR knockdown.

### Prolonged loss of TPR during senescence blocks NF-κB activation

Because of the enrichment for NF-κB motifs in the d8 SEN^+^TPR^+^ putative enhancers, we set out to investigate whether NF-κB activation is affected by TPR knockdown in senescent cells.

Inactive NF-κB dimers are held in the cytoplasm through their association with IκB proteins. Inducing stimuli trigger activation of the IκB kinase complex (IKK), which leads to phosphorylation and degradation of IκB, allowing the translocation of NF-κB to the nucleus, where it promotes the transcription of target genes (Hayden and Ghosh, 2012). We used immunofluorescence to assess NF-κB localisation in the nucleus and cytoplasm during OIS and in the presence or absence (siRNA knockdown) of TPR (Figure 2A). As expected, NF-κB remained cytoplasmic in control (STOP) cells, but translocation to the nucleus could be detected in senescent RAS cells, with SAHF readily apparent from DAPI staining in the nucleus of these cells. As we previously reported, knockdown of TPR (siTPR) in RAS cells blocks SAHF formation, but it also results in reduced nuclear localisation (decreased nucleocytoplasmic ratio) of NF-κB, consistent with decreased NF-κB activation (Figure 2A and B, Figure S2A).

**Figure 2.**
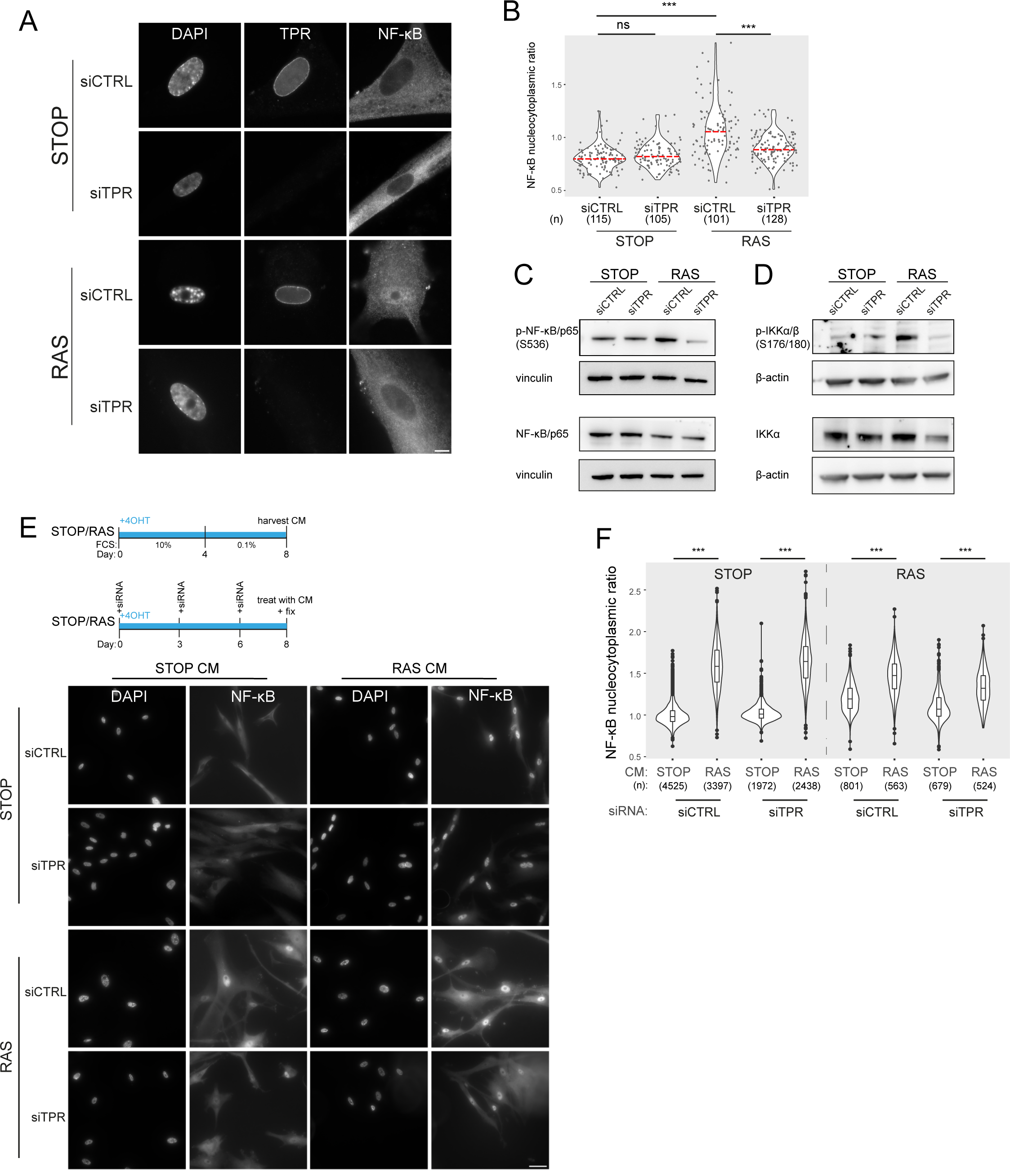
Prolonged loss of TPR during senescence blocks NF-ĸB activation. A) TPR and NF-ĸB immunostaining in control (STOP) and OIS (RAS) cells after 4-OHT and siRNA (control and TPR) treatment for 8 days. Scale bar: 10 µm. B) Quantification of NF-ĸB nucleocytoplasmic ratios in experiment described in (A). Kruskal-Wallis testing was used to determine statistical significance followed by Dunn post-hoc testing, n.s. p >0.05, *** < 0.001. (n) indicates the number of cells analysed for each sample. Data from a biological replicate are in Figure S2A. C) Immunoblots of extracts from control (STOP) and OIS (RAS) cells after 4-OHT and siRNA treatment for 8 days for phosphorylated (pS536) and total NF-kB with vinculin as a loading control. D) As in (C) but for phosphorylated (pS176/180) IKKα/β and total IKKα and with β-actin as a loading control. Data from biological replicates of (C) and (D) are in Figures S2B and C. E) Left: Schematic of controlled media experiment to investigate whether TPR loss causes a general defect in NF-ĸB transport. STOP and RAS cells were grown for 8 days and treated with 4-OHT and siRNAs. On d8 they were treated for 45 minutes with conditioned media (CM) taken from either STOP or RAS cells grown in 4-OHT-containing media for 8 days. Right: NF-ĸB immunostaining in STOP or RAS cells treated with CM harvested from STOP (left) or RAS (right) cells. Scale bar: 50 µm. F) Quantification of NF-ĸB nucleocytoplasmic ratios for experiment shown in (E). Data from a biological replicate are in Figure S2D.

**Figure S2.**
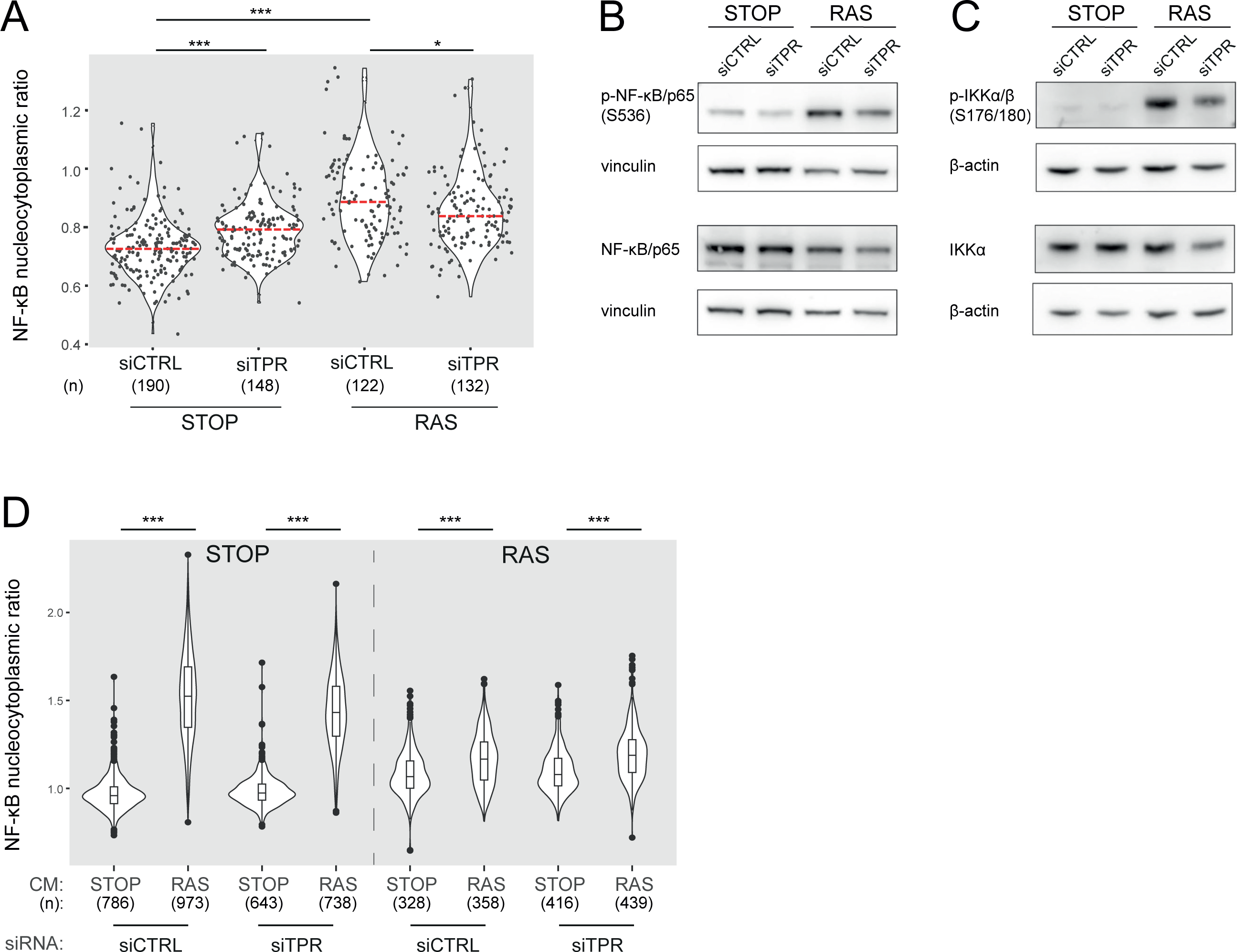
TPR depletion blocks NF-kB activation during senescence. Related to Figure 2. A) Quantification of NF-ĸB nucleocytoplasmic ratios by immunofluorescence in STOP and RAS cells after 4-OHT and siRNA treatment for 8 days. Kruskal-Wallis testing was used to determine statistical significance followed by Dunn post-hoc testing, n.s. p >0.05, *** < 0.001. (n) indicates the number of cells analysed for each sample. Data are from a biological replicate of the experiment shown in Figure 2B. B) Immunoblots in extracts from control (STOP) and OIS (RAS) cells after 4-OHT and siRNA (control and TPR) treatment for 8 days for phosphorylated (pS536) and total NF-ĸB with vinculin as a loading control. Biological replicate of data in Figure 2C. C) As in (B) but for phosphorylated (pS176/180) IKKα/β and total IKKα and with β-actin as a loading control. Biological replicate of data in Figure 2D. D) Quantification of NF-ĸB nucleocytoplasmic ratios in d8 STOP and RAS treated for 45 minutes with conditioned media (CM) from either STOP or RAS cells grown in 4-OHT-containing media for 8 days. Data are from a biological replicate of experiment in Figure 2F.

Active NF-κB is phosphorylated at serine 536 (Sakurai et al., 1999). Immunoblotting showed that phosphorylation of NF-κB decreased upon TPR knockdown in senescent cells (Figure 2C, Figure S2B). Phosphorylation of the NF-κB kinase IKK was reduced upon TPR knockdown (Figure 2D, Figure S2C), further suggesting a reduction in NF-κB signalling pathway activation in senescent cells in the absence of TPR.

As TPR is part of the nuclear pore basket, it is possible that the knockdown of TPR causes a general defect in nuclear import, preventing activated NF-κB being imported into the nucleus upon OIS. To check that this is not the case, we grew control and RAS senescent cells, treated with 4-OHT and siRNAs as before, then exposed them to conditioned media (CM) from either control or RAS cells enriched in SASP factors (8 days post 4-OHT treatment) (Figure 2E). Conditioned media from senescent cells is enriched in SASP factors which leads to NF-κB activation (Boumendil et al., 2019). Immunofluorescence showed that NF-κB nuclear translocation occurred in RAS cells (with control siRNA) in the presence of CM from either STOP or RAS cells, because of their intrinsic activation of the SASP. In STOP cells, nuclear translocation of NF-κB was only induced by CM from RAS cells. This was not affected by TPR knockdown, and this was also the case for RAS cells after TPR knockdown (Figure 2E). Quantification of the NF-κB nucleocytoplasmic ratio confirmed that TPR knockdown does not affect the nuclear import of NF-κB (Figure 2F, Figure S2D).

### Decreased NF-κB activation upon TPR knockdown precedes the SASP

The SASP reinforces itself via a positive feedback loop – once secreted, SASP factors bind to their receptors on the cell membrane, leading to NF-κB activation and increased SASP (Figure 3A) (Acosta et al., 2008; Freund et al., 2010; Orjalo et al., 2009). Therefore, decreased NF-κB activation at d8 of RAS induction upon TPR knockdown could be result from a general decrease in the SASP. To determine whether this was the case, we assessed NF-κB nuclear localisation at two earlier timepoints: day 3 (d3), which is before SASP initiation and occurs when the cells are coming out of the initial highly proliferative state (Young et al., 2009); and day 5 (d5), which is at the initial stages of the inflammatory SASP (Figure 3B). There was no change in NF-κB nucleocytoplasmic ratio at d5 between any of the samples, and only a small increase between STOP siCTRL and RAS siCTRL at d3 (Figure 3C, Figure S3A), suggesting that these timepoints may be too early to observe significant NF-κB nuclear translocation. However, nuclear NF-κB intensity in the cell was increased in OIS-induced RAS cells compared with the control STOP cells at both d3 and d5, suggesting early NF-κB activation (Figure 3D, Figure S3B). Knockdown of TPR led to significantly lower nuclear NF-κB intensities in RAS cells at both timepoints, suggesting early NF-κB signalling is reduced when OIS is induced in the absence of TPR. A small increase in NF-κB nuclear intensity in d3 STOP cells when TPR was knocked down and a small decrease at d5 were not reproducible (Figure 3D, Figure S3B). Consistent with an effect on early NF-κB activation, immunoblotting showed that TPR knockdown resulted in decreased NF-κB phosphorylation (S536) in RAS cells at both d3 and d5 (Figure 3E, Figure S3C), and decreased phosphorylation of IKK, the upstream kinase (Figure 3F, Figure S3D).

**Figure 3.**
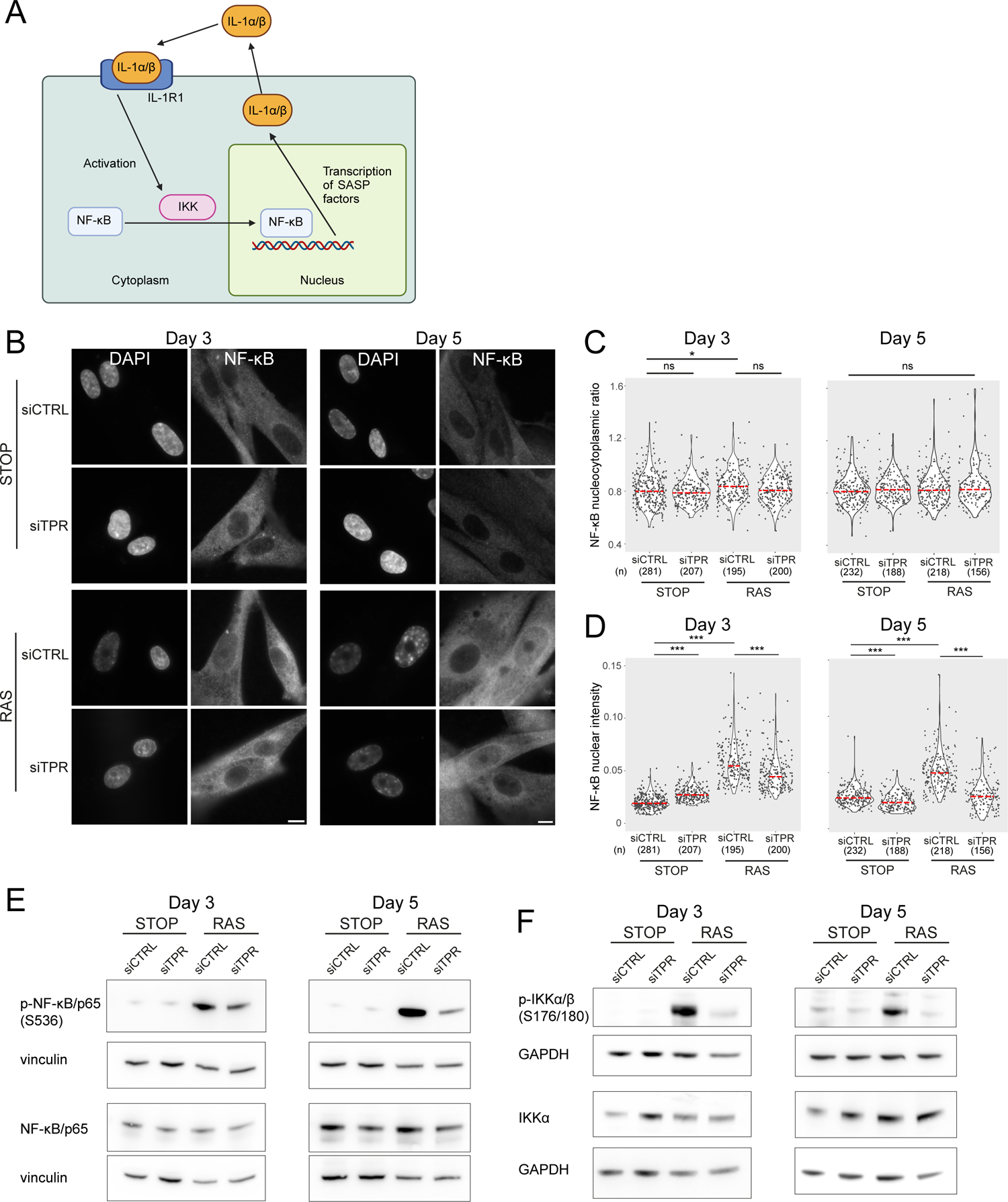
Decreased NF-ĸB activation upon TPR knockdown precedes the SASP. A) Schematic showing positive feedback loop in SASP signalling. Secreted IL-1α and IL-1 β bind IL-1R1 at the cell membrane, leading to increased NF-ĸB activation and increased IL-1α and IL-1 β secretion. B) NF-ĸB immunostaining in control (STOP) and OIS (RAS) cells after 4-OHT and siRNA treatment for either 3 or 5 days. Scale bar: 10 µm. C and D) Quantification of (C) nucleocytoplasmic ratios of NF-ĸB or (D) NF-ĸB nuclear intensity from experiment shown in (B). (n) indicates the number of cells analysed for each sample. Kruskal-Wallis testing was used to determine statistical significance followed by Dunn post-hoc testing, n.s. p > 0.05, * < 0.05, *** < 0.001. E) Immunoblots for phosphorylated (pS536) and total NF-ĸB (p65) in STOP and RAS cells treated with 4-OHT for 3 or 5 days and with control (CTRL) or TPR siRNAs. Vinculin was used as a loading control. F) As in (E) but blotting to detect phosphorylated (pS176/180) IKKα/β and total IKKα. GAPDH was used as a loading control. Data from a biological replicate of the data in Figure 3 are in Figure S3.

**Figure S3.**
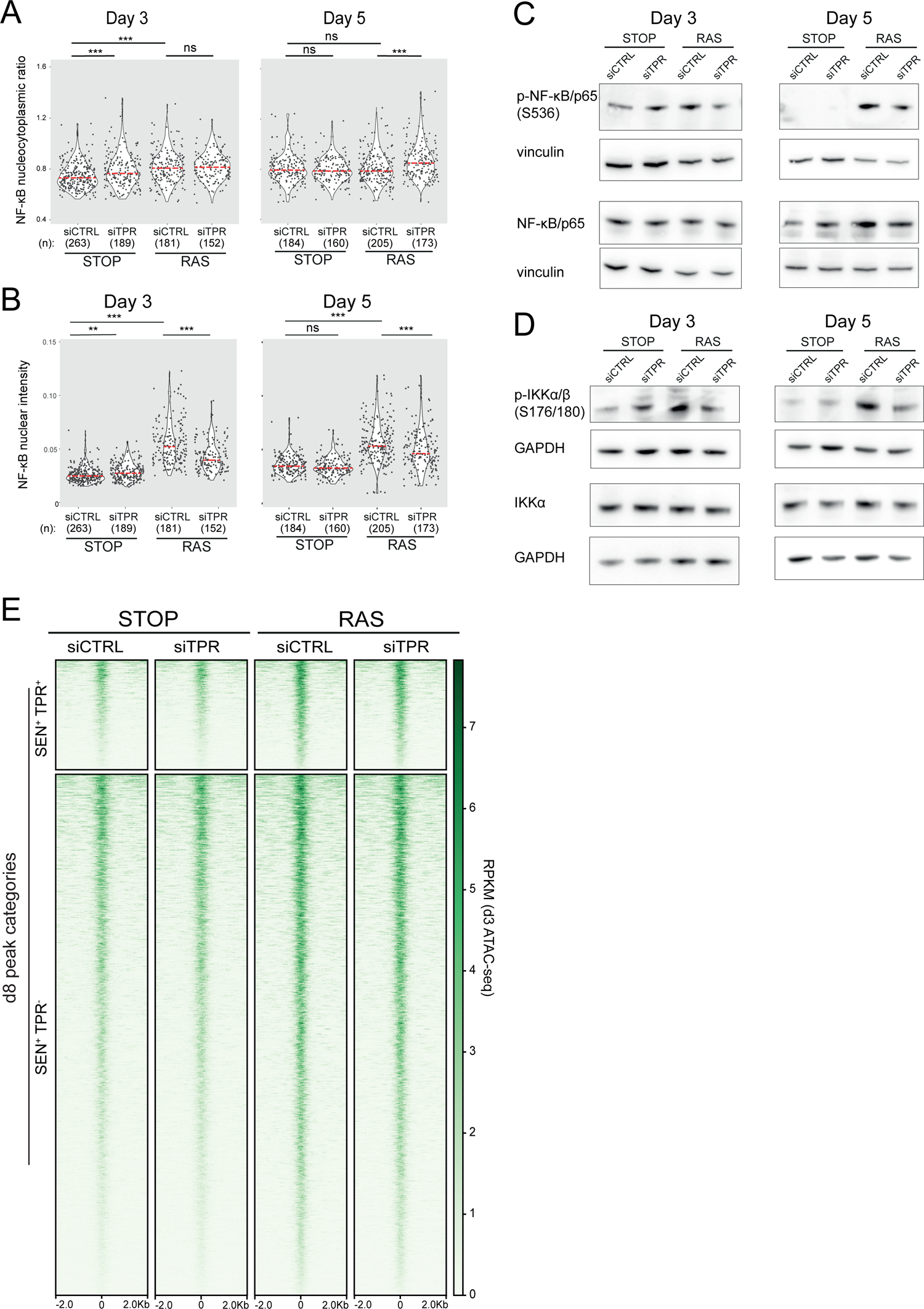
Decreased NF-ĸB activation upon TPR knockdown at days 3 and 5. Related to Figure 3. A and B) Quantification of (A) nucleocytoplasmic ratios of NF-kB or (B) nuclear NF-ĸB intensity from a biological replicate of the experiment shown in Figure 3B-D. (n) indicates the number of cells analysed for each sample. Kruskal-Wallis testing was used to determine statistical significance followed by Dunn post-hoc testing, n.s. p> 0.05, * < 0.05, *** < 0.001. C) Immunoblots for phosphorylated (pS536) and total NF-ĸB (p65) in STOP and RAS cells treated with 4-OHT for 3 or 5 days and with control (CTRL) or TPR siRNAs. Vinculin was used as a loading control. Biological replicate of data in Figure 3E. D) As in (C) but blotting to detect phosphorylated (pS176/180) IKKα/β and total IKKα. GAPDH was used as a loading control. Biological replicate of data in Figure 3F. E) Heatmap showing ATAC-seq signal in control (STOP) and OIS (RAS) cells 3 days after 4-OHT treatment and transfected with either CTRL or TPR siRNAs. Peak categories are those defined from the d8 ATAC-seq data in Figure 1C. Intensity scale represents RPKM.

To determine whether, as at d8, lowered NF-κB activity upon TPR knockdown during early RAS activation (day 3) is accompanied by changes in chromatin accessibility at the enhancers of SASP genes, we carried out ATAC-seq on STOP and RAS cells treated with 4-OHT for 3 days, as well as with control and TPR siRNAs. Whilst some of the regions defined as senescence specific (SEN^+^) at d8 also show senescence-specific enhanced chromatin accessibility at d3, albeit less marked than at d8, SEN^+^ accessibility peaks that were TPR-dependent (TPR^+^) at d8 did not show decreased chromatin accessibility upon TPR knockdown at d3 (Figure S3E). Indeed, TPR knockdown did not lead to any significant changes in ATAC-seq peaks in either STOP or RAS cells (Table S2). Therefore, TPR plays a role in NF-κB activation during the early stages of stress in response to oncogenic RAS, before activation of the SASP and without affecting chromatin accessibility at enhancers.

### TPR knockdown during the early stages of OIS reduces STING expression and TBK1 activation in response to the stress induced by oncogenic RAS

Although we detect no changes in chromatin accessibility upon TPR knockdown at day 3 of oncogenic stress, the decrease in NF-κB activation suggests that the initial signalling events leading to the loss of the SASP are already occurring. We therefore used RNA sequencing (RNA-seq) to investigate the transcriptional changes that could be driving the TPR-dependent decrease in NF-κB activation at d3.

Through its interaction with the TREX-2 complex, TPR is known to be required for the specific export of intronless and intron-poor mRNAs, as well as histone mRNAs, the majority of which are intronless (Aksenova et al., 2020; Lee et al., 2020). Indeed, of the genes downregulated on TPR knockdown, 14% (STOP) or 13% (RAS) are intronless (Fisher’s exact test, p = 1.2 x 10^-11^; p = 7.1 x 10^-9^, respectively) (Figure S4A). This includes histone genes (STOP cells: 3 genes, p = 6.1 x 10^-3^; RAS cells: 6 genes, p = 4.2 x 10^-6^) (Figure S4B).

**Figure S4.**
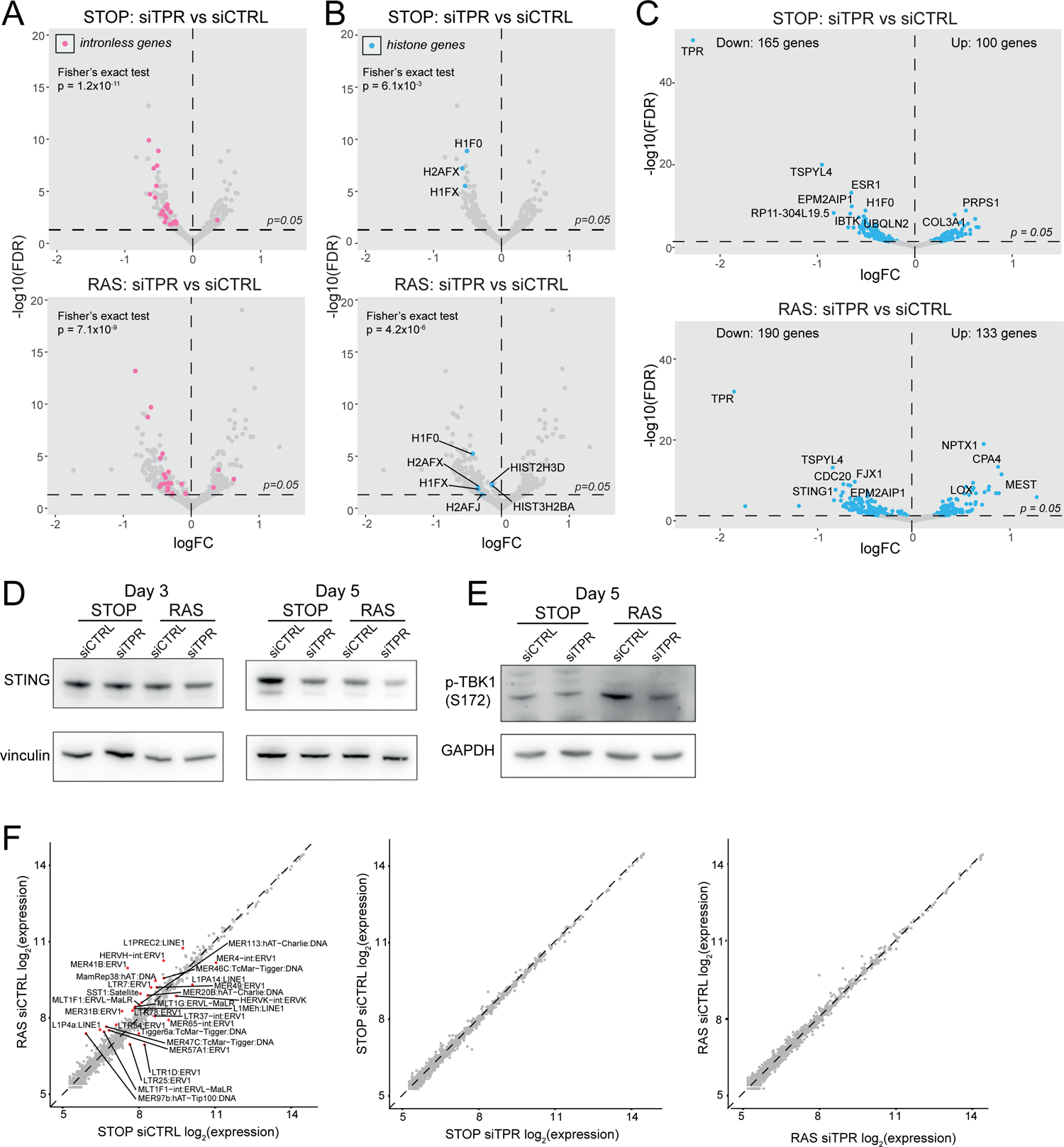
Decreased abundance of mRNAs for intronless genes and for *STING1* in RAS cells upon TPR knockdown at day 3. Related to Figure 4. A) Volcano plots of differential expression analysis of d3 STOP (top) and RAS (bottom) cells treated with TPR vs CTRL siRNAs with intronless genes coloured pink. Horizontal dashed line indicates an adjusted p-value (FDR) of 0.05. Axes are truncated for clarity so change in expression for TPR is not shown. Fisher’s exact test (p) was carried out to determine whether the number of downregulated intronless genes was greater than expected by chance. (B) As in (A) but with histone genes labelled in blue. C) Volcano plot showing differential expression analysis comparing siTPR vs siCTRL in d3 STOP (top) or RAS (bottom) cells. Blue dots indicate differentially expressed genes and the dashed horizontal line indicates an adjusted p-value of 0.05. The 10 genes with the most significant p-values are labelled. D) Immunoblots detecting STING in STOP and RAS cells treated with 4-OHT for 3 or 5 days and with control (siCTRL) or TPR siRNAs. Vinculin was used as a loading control. Biological replicate of the data in Figure 4C. E) As in (D) but detecting phosphorylated TBK1 (pS172) in STOP and RAS cells at d5 of OIS. GAPDH was used as a loading control. Biological replicate of the data in Figure 4D. G) Log-transformed RNA-seq counts for RNAs transcribed from transposable elements comparing (top) RAS vs STOP cells at d3 and treated with control siRNAs; (middle) d3 STOP cells treated with control vs TPR siRNAs; (bottom) d3 RAS cells treated with control vs TPR siRNAs. Transposable elements with absolute fold changes > 1.5 and adjusted p-values < 0.05 are highlighted in red and labelled.

*TPR* was the most significantly downregulated gene when comparing siTPR with siCTRL in both RAS and STOP cells (Figure S4C). To determine which changes in expression were specific to cells undergoing oncogenic stress, we compared RAS siTPR with RAS siCTRL, disregarding any genes that also changed in expression upon TPR knockdown in STOP cells. *STING1* showed the most significant RAS-specific decrease in expression (Figure 4A). Reduced *STING1* mRNA in RAS cells after 3 days of RAS induction and TPR knockdown was validated by RT-qPCR (Figure 4B). Immunoblotting did not detect a reduction in levels of STING protein at d3 of RAS induction, perhaps due to protein stability at this short time point, but decreased levels were detected by d5 (Figure 4C and Figure S4D).

**Figure 4.**
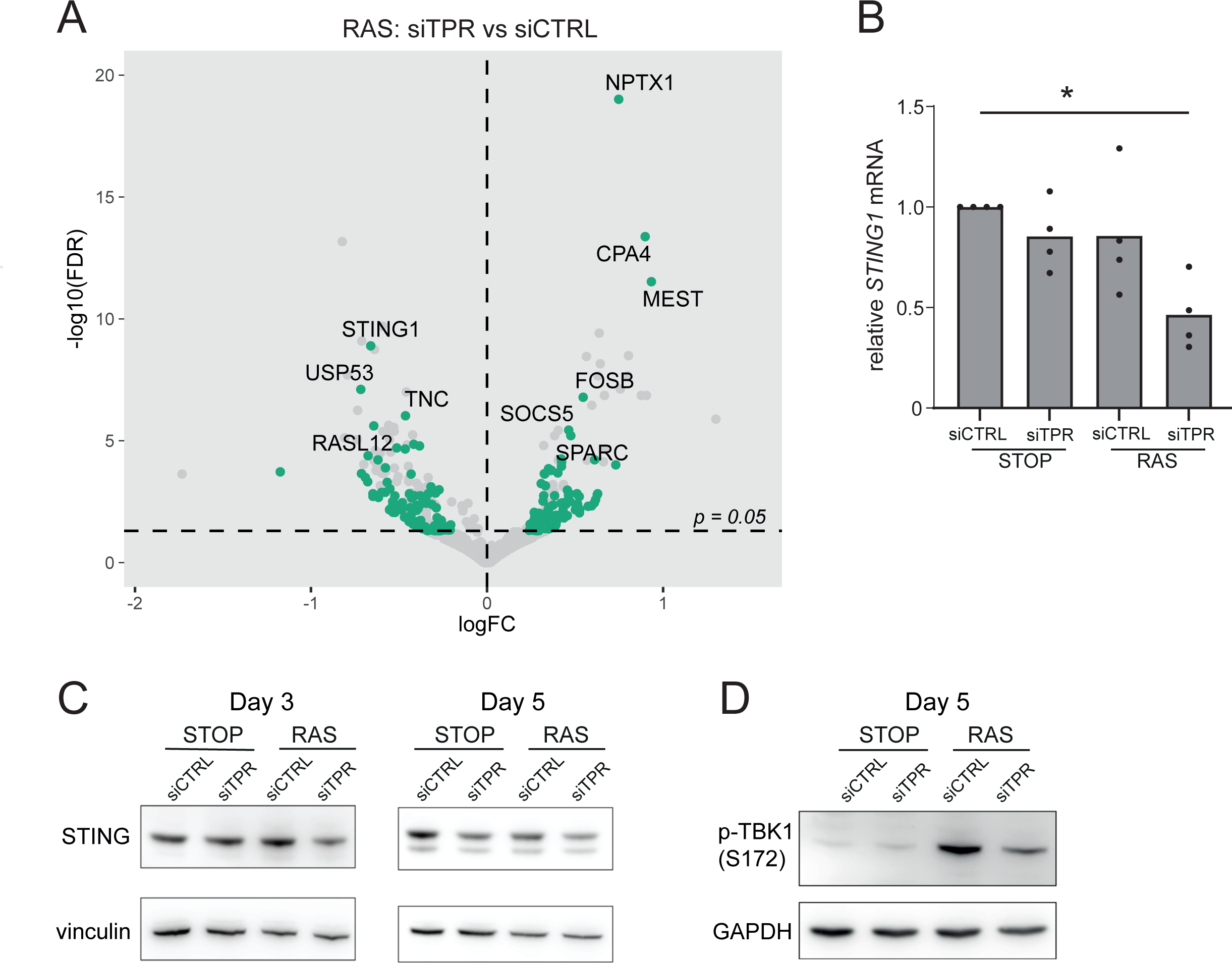
Decreased STING expression and TBK1 activation upon TPR knockdown during early stages of OIS. A) Volcano plot of differential expression analysis of RNA isolated from RAS cells at d3 of OIS and treated with siTPR vs siCTRL. Genes showing a significant change in expression in RAS, but not in STOP cells are indicated in green and the 10 most significant of these are labelled. Horizontal dashed line indicates an adjusted p-value (FDR) of 0.05. Axes are truncated for clarity so the change in expression for TPR is not shown. B) RT-qPCR for *STING1* mRNA in RNA prepared from STOP and RAS cells treated with 4-OHT for 3 days and with control (siCTRL) and TPR siRNAs. Expression is relative to STOP cells treated with siCTRL and normalised to levels of *GAPDH* mRNA. Individual data points are the mean of three technical replicates for each of four biological replicates. An ANOVA test was used to determine statistical significance. * < 0.05. C) Immunoblots detecting STING in STOP and RAS cells treated with 4-OHT for 3 or 5 days and with control (siCTRL) or TPR siRNAs. Vinculin was used as a loading control. D) As in (C) but detecting phosphorylated TBK1 (pS172) in STOP and RAS cells at d5 of OIS. GAPDH was used as a loading control. Data from biological replicates for (C) and (D) are in Figure S4C and D.

The cGAS-STING pathway is known to activate the SASP via NF-κB signalling (Dou et al., 2017; Glück et al., 2017; Yang et al., 2017). TANK-binding kinase 1 (TBK1) acts downstream of STING-mediated sensing of cytosolic DNA, and controls NF-κB signalling. TBK1 is phosphorylated at serine 172 when active (Abe and Barber, 2014; Shu et al., 2013). To investigate whether TPR is required for activation of this pathway early in OIS, we therefore analysed TBK1 phosphorylation. Immunoblotting showed decreased TBK1 phosphorylation in RAS cells upon TPR knockdown at d5 of RAS induction (Figure 4D and Figure S4E).

The transcription of several classes of retrotransposons, including long-interspersed element-1 (LINE1) and human endogenous retroviruses (HERVs), is known to be activated in senescent cells, and sensed through cGAS triggering an innate immune response (De Cecco et al., 2019; Liu et al., 2023). To investigate whether TPR-dependent regulation of cGAS-STING/TBK1 signalling could act through an effect on retrotransposon expression, we used TEtranscripts (Jin et al., 2015) to quantify the abundance of RNAs from transposable elements in d3 of RAS induction RNA-seq data. Although RNA abundance for some transposable elements, including HERV and LINE1 elements, was higher in RAS compared with STOP cells treated with control siRNA, there were no significant changes in transposable element RNA abundance upon knockdown of TPR in either cell line (Figure S4F). This suggests that it is not a change in transposable element expression that drives the decrease in innate immune signalling seen upon TPR knockdown at d3 of OIS.

### TPR is required for the formation of cytoplasmic chromatin fragments during the early stages of OIS

Another trigger of innate immune activation in senescent cells is the generation of CCFs (Dou et al., 2017; Glück et al., 2017; Yang et al., 2017). To determine whether TPR affects the generation of CCFs, we assessed their frequency – as evidenced by the proportion of cells with DAPI-stained foci in the cytoplasm not obviously connected to the nucleus at d3 and d5 of RAS induction. The frequency of detectable CCFs decreased when TPR was knocked down. Though apparent by d3, this was only statistically significant at d5 (Figure 5A).

**Figure 5.**
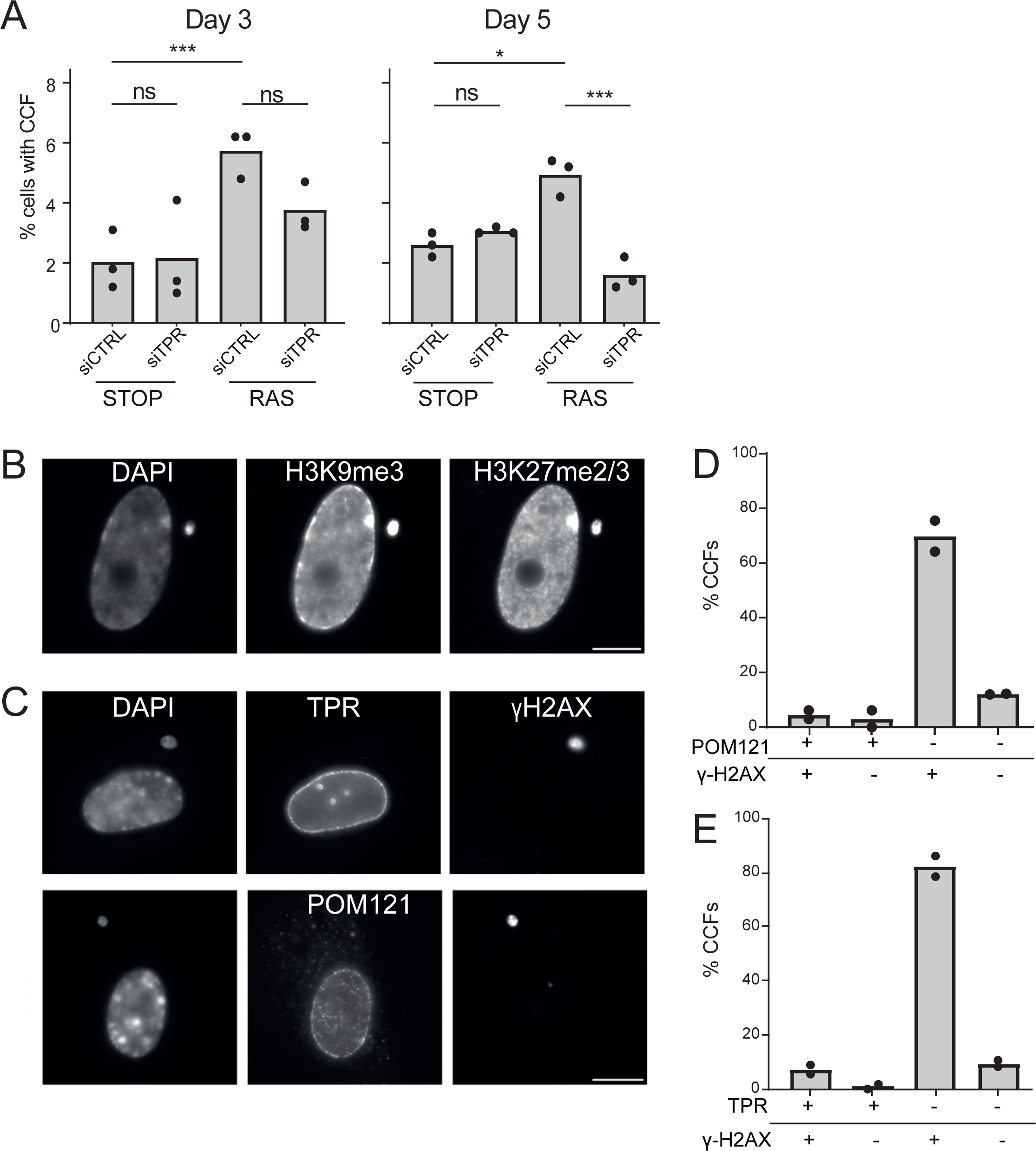
TPR is required for the induction of CCFs during the early phase of OIS. A) Mean percentage of cells containing CCFs in STOP and RAS cells at d3 or d5 of OIS and treated with either control (siCTRL) or TPR siRNAs. Data points are for three biological replicates. Data were fitted to a generalised linear model before carrying out pairwise comparisons between samples, n.s. p > 0.05, * < 0.05, *** < 0.001. B) Immunostaining for H3K9me3 and H3K27me2/3 in a DAPI-stained d5 RAS cell with a CCF. Scale bar: 10 µm. C) As in (B) but in d5 RAS cells containing CCFs and staining for γH2AX and either TPR (top) or POM121 (bottom). Scale bar: 10 µm. D) Mean percentage of CCFs that show +ve or -ve staining for POM121 or γ-H2AX in d5 RAS cells. Data are from two biological replicates (n = 49 and 67 CCFs). E) Mean percentage of CCFs that show +ve or -ve staining for TPR or γ-H2AX in d5 RAS cells. Data are from two biological replicates (n = 56 and 36 CCFs).

CCFs form from blebbing off of the nuclear membrane, thought to be as a result of a loss of structural integrity of the nuclear envelope (Ivanov et al., 2013). CCFs are known to contain lamin B1 (Dou et al., 2015) but whether they contain other components of the nuclear envelope is unexplored. By immunostaining we confirmed that, as expected, CCFs are positive for the heterochromatic histone marks H3K9me3 and H3K27me3 (Figure 5B) and for γ-H2AX (Dou et al., 2017, 2015). However, they appear to lack staining for TPR or for POM121, a transmembrane nucleoporin, suggesting that there are no nuclear pores in the CCF envelope (Figure 5C-E).

## Discussion

We have previously linked TPR at the nuclear pore basket to the nuclear reorganisation of heterochromatin away from the nuclear periphery to form senescence-associated heterochromatic foci (SAHF), and to the activation of SASP genes, during the process of oncogene-induced senescence (Boumendil et al., 2019). The extent to which these events are coupled was unclear. In this study, we address this by looking at the effects of depleting TPR knockdown very early (day 3) following the induction of oncogenic RAS, as the cells are responding to the initial stress and before SASP gene transcriptional induction (Young et al., 2009).

We show that TPR loss during OIS (d8) prevents the chromatin opening of SASP gene enhancers enriched in binding motifs for NF-κB - a key transcription factor that drives the SASP. However, we show that TPR is also required for the very early stages of NF-κB activation upon RAS oncogenic stress, well before SASP gene activation, suggesting that TPR does not have a direct effect on chromatin structure at enhancers of the SASP. Rather, our data suggest that TPR loss has its impact upstream of NF-κB and its translocation to the nucleus, by decreasing TBK1 phosphorylation, likely downstream of cGAS-STING signalling. We link this to the production of cytoplasmic chromatin fragments (CCFs) – the number of CCFs decreases when TPR is knocked down. This is intriguing, due to the proximity of TPR to the nuclear envelope, which is where CCFs are generated.

Cytoplasmic chromatin derived from the nuclear genome is known to activate the innate immune response, sensed and signalled through cGAS-STING upstream of TBK1. Cytoplasmic DNA sensing is best studied in the context of micronuclei, formed during mitosis as a consequence of unrepaired DNA damage (Miller et al., 2021). Micronuclei can contain many different types of chromatin (Mammel et al., 2021) and have been reported to have NPCs in their membrane, albeit at a much lower density than the primary nuclear membrane (Crasta et al., 2012; Hatch et al., 2013; Liu et al., 2018). In contrast, we do not detect a core nuclear pore component (POM121), or TPR, in CCFs, consistent with a mechanism of formation that is distinct from that of micronuclei. This suggests that either the CCFs are formed from the nuclear membrane between NPCs, or that NPCs are rapidly lost from CCFs.

CCFs form by blebbing off from the nuclear periphery (Ivanov et al., 2013; Miller et al., 2021) and preferentially contain chromatin fragments enriched in the repressive histone modifications (H3K9me3 and H3K27me3) that are typically located at the nuclear periphery in lamina associated domains (Guelen et al., 2008). Since we have shown that TPR depletion prevents the loss of heterochromatin from the nuclear periphery and the formation of SAHF during OIS, it is plausible that the decrease in CCFs produced during the early phases of OIS upon TPR knockdown may be caused by an increase in the stability of the nuclear periphery due to the heterochromatin that remains there when SAHF are not formed.

TPR has also been suggested to interact with, and affect the organisation of, lamin B1 at the nuclear periphery (Fišerová et al., 2019) and there is loss of lamin B1 from the nuclear periphery in senescence (Dou et al., 2015; Freund et al., 2012; Sadaie et al., 2013; Shimi et al., 2011). Therefore, it is possible that TPR depletion also impacts CCF formation through its effects on lamin B1. Finally, TPR may be involved in bending of the nuclear membrane during NPC assembly in interphase (Otsuka et al., 2023), so another possibility is that TPR contributes to nuclear membrane curvature, perhaps enabling nuclear membrane blebbing during CCF formation. Our results suggest a role for TPR as an important factor in the loss of nuclear integrity that occurs in response to oncogene-induced stress, leading directly to activation of cytoplasmic nucleic acid sensing and the key inflammatory gene expression programme of senescence. Whether TPR has a similar role for other triggers of senescence and in aging remains to be determined.

## Methods

### Cell culture, conditioned media preparation and siRNA transfection

IMR90 cells were cultured in DMEM with 10% FBS and 1% penicillin/streptomycin in a 37°C incubator with 5% CO_2_. IMR90 cells were infected with pLNC-ER:RAS and pLXS-ER:STOP retroviral vectors to produce RAS and STOP cells respectively (Acosta et al., 2013). RAS translocation to the nucleus was induced by addition of 100 nM 4-hydroxytamoxifen (4-OHT) (Sigma). 4-OHT-containing medium was changed every 3 days.

To prepare conditioned media, 5 x 10^5^ STOP and RAS cells were grown in 100 nM 4-OHT media for 8 days. After 4 days this was replaced with media with 0.1% FCS and 100 nM 4-OHT. Media was harvested on day 8. To activate NF-κB, cells were treated with the conditioned media for 45 minutes (mins).

siRNA knockdown was carried out as previously described (Boumendil et al., 2019). Briefly, 9 x 10^5^ STOP or RAS IMR90 cells (except for imaging experiments, which used 1.5 x 10^5^ cells) were transfected using Dharmafect transfection reagent (Dharmacon) with a 30 nM final concentration of control (siCTRL, D-001810-10-59) or TPR (siTPR, L-010548-00) siRNA pools (Dharmacon). Transfections were carried out on day 0 of 4-OHT treatment and on every third subsequent day.

### Immunofluorescence

Cells were seeded onto coverslips 48 hours before fixation with 4% paraformaldehyde in PBS for 30 mins at room temperature, before permeabilization with 0.2% Triton X-100 for 10 mins. Coverslips were then washed three times with PBS before blocking in 1% bovine serum albumin (BSA) for 30 mins. Coverslips were then incubated for 45 mins in a humid chamber with primary antibody diluted in 1% BSA at the dilutions detailed in Table S3. Coverslips were washed three times with PBS. Cells were then incubated with fluorescently labelled secondary antibodies (Life Technologies, Table S3) for 30 mins followed by two washes in PBS. Finally, PBS with 50 ng/ml DAPI was added for 4 mins, before a final wash with PBS and mounting onto slides with VectaShield (Vector Laboratories).

Epifluorescence images were acquired using either a Photometrics Coolsnap HQ2 CCD camera (Teledyne Photometrics) or a Hamamatsu Orca Flash 4.0 CMOS camera on a Zeiss Axioplan II fluorescence microscope with Plan-neofluar/apochromat objective lenses (Carl Zeiss UK), a Mercury Halide fluorescent light source (Exfo Excite 120, Excelitas Technologies) and Chroma #83000 triple band pass filter set (Chroma Technology Corp.) with the single excitation and emission filters installed in motorised filter wheels (Prior Scientific Instruments). Image capture was performed using Micromanager (Version 1.4).

For the conditioned media experiment (Figure 2) images were acquired using a Photometrics Prime BSI CMOS camera (Teledyne Photometrics) fitted to a Zeiss AxioImager M2 fluorescence microscope with Plan-Apochromat objectives, a Zeiss Colibri 7 LED light source, together with Zeiss filter sets 90 HE, 92 HE, 96 HE, 38 HE and 43 HE (Carl Zeiss UK). Image capture was performed in Zeiss Zen 3.5 software.

### Image analysis

Nuclear NF-κB intensity and nucleocytoplasmic ratios were calculated using CellProfiler (Stirling et al., 2021). Nuclei were identified in the DAPI channel using the Identify Primary Objects module to carry out adaptive Otsu thresholding with a threshold smoothing scale of 5, a threshold correction factor of 0.37, a 200-pixel adaptive window and a typical object diameter of 100-500 pixels. Clumped objects were distinguished using the ‘Intensity’ method and dividing lines were drawn between clumped objects using the ‘Shape’ method. A secondary object was then generated by expanding the primary object by 50 pixels, and NF-κB intensity measured for the primary object (nucleus), secondary object (cell), and a tertiary object (cytoplasm) generated by removing the primary object area from the secondary object. Nucleocytoplasmic ratio was calculated by dividing the NF-κB intensity in the cytoplasm by the nuclear NF-κB intensity.

To count cytoplasmic chromatin fragments (CCFs), 500 cells per sample were observed by epifluorescence microscopy and cells displaying cytoplasmic DAPI staining were imaged. One blinded replicate was carried out by S.B. who was unfamiliar with previous results. For quantification of CCFs with γ-H2AX, TPR and POM121 staining, all cells on a slide of d5 OIS RAS cells were assessed and imaged if they displayed cytoplasmic DAPI staining.

### Immunoblotting

Cells were lysed in Cell Lysis Buffer (20 mM Tris-HCl pH 7.5, 150 mM NaCl, 1 mM Na_2_EDTA, 1 mM EGTA, 1% Triton-X100, 2.5 mM sodium pyrophosphate, 1 mM β-glycerophosphate, 1 mM Na_3_VO_4_, 1 μg/ml leupeptin, Cell Signaling Technology) with one Pierce Phosphatase and Protease Inhibitor Mini Tablet (Thermo Fisher) added per ml of cell lysate. Protein concentration was quantified using a Pierce BCA protein analysis kit (Thermo Fisher), and then 20μg of protein was run on NuPage 4-12% Bis-Tris gels (Thermo Fisher) at 150V for 1 hour. After transfer onto nitrocellulose membranes with an iBlot 2 gel transfer device (Thermo Fisher), membranes were blocked in 5% BSA in TBS with 0.1% Tween-20 (TBS-T) for 30 mins then incubated overnight with the primary antibodies at the dilutions detailed in Table S3, in 5% BSA in TBS-T. After 3 x 10 min washes in TBS-T, membranes were incubated with the appropriate horseradish peroxidase (HRP)-conjugated secondary antibodies, before three further washes with TBS-T. Membranes were imaged using an Amersham ImageQuant 800 imager (Cytiva) on the chemiluminescence setting with the SuperSignal West Femto maximum sensitivity substrate kit (Thermo Fisher). When using the mouse anti-β-actin−HRP antibody, the primary antibody incubation step was omitted and a 10 min incubation carried out alongside the secondary antibody step for other blots, before washing and imaging as before.

### ATAC-seq library preparation

A standard ATAC-seq protocol with IMR90 cells yielded too many mitochondrial reads and high PCR duplication levels because of poor tagmentation. To circumvent this issue, we used the Omni-ATAC protocol (Corces et al., 2017) with some modifications. Briefly, IMR90 cells were harvested by trypsinization and washed with cold PBS. One million cells were resuspended in ice cold ATAC resuspension buffer (ATAC-RSB; 20 mM Tris HCl pH 7.6, 10 mM MgCl_2_, 20% dimethyl formamide) and 40 strokes in a 1 ml Dounce using a rounded pestle were applied. Debris was pre-cleared by spinning at 100 g for 3 mins. The supernatant was collected and spun again at 1000 g for 5 min to collect the nuclear pellet. The pellet was resuspended in 1 ml ATAC-RSB buffer with 0.1% Tween-20 and spun at 1000 g for 5 min. The nuclear pellet was resuspended in 100 μl TD buffer (10 mM Tris HCl pH 7.6, 5 mM MgCl_2_, 10% dimethyl formamide) and the Omni-ATAC protocol performed on 5×10^4^ nuclei. ATAC-seq libraries were made using adaptor sequences as described previously (Buenrostro et al., 2013). Libraries were assessed for quality and fragment size using the Agilent Bioanalyser. Sequencing was performed on the NextSeq 2000 platform (Illumina) using NextSeq 1000/2000 P2 Reagents.

### ATAC-seq data analysis

FastQC was used to obtain basic quality control metrics from sequencing data and to assess the quality of reads before preprocessing steps. Sequencing reads were trimmed to a minimum of 30 bases and adaptor sequences clipped using cutadapt (Martin, 2011). Reads were aligned to the human genome assembly hg19 using bowtie2 (Langmead and Salzberg, 2012). Mitochondrial reads and PCR duplicates were filtered out before shifting reads by +4 bp for the positive strand and −5 bp for the negative strand. Peaks were then called using MACS2 (Zhang et al., 2008) before removing all peaks from promoter regions, as we were specifically interested in promoter-distal regulatory elements. The HOMER (Heinz et al., 2010) functions makeTagDirectory and annotatePeaks.pl with settings ‘-noadj -len 0 -size given’ were used for read counting before count tables were loaded into RStudio.

Trimmed Mean of M-values (TMM) normalisation was carried out using edgeR (Robinson et al., 2010) and analysis of differentially accessible regions was carried out using limma (Ritchie et al., 2015). Contrasts were designed as ∼0+Sample, where Sample specifies both the cell line and siRNA treatment. A cut-off adjusted p-value of 0.05 was used to define differentially accessible peaks. Heatmaps were generated using the deepTools function plotHeatmap (Ramírez et al., 2016). Analysis of nearby genes was carried out using GREAT (McLean et al., 2010) with the ‘basal plus extension’ setting. Motif analysis was carried out using HOMER (Heinz et al., 2010).

### Analysis of published ChIP-seq data

NarrowPeak files for H3K27 acetylation ChIP-seq from growing and senescent IMR90 RAS^G12V^ cells (Parry et al., 2018) were obtained from the Gene Expression Omnibus with accession number GSE103590. Correlation between replicates was checked using the plotCorrelation function from the deepTools package (Ramírez et al., 2016). Heatmaps were generated by using the deepTools function plotHeatmap to plot the first replicate from each sample with peak categories taken from the ATAC-seq analysis.

### RT-qPCR

Total RNA was extracted using the RNeasy mini kit (Qiagen) and cDNAs generated using SuperScript II (Life Technologies). Real-time PCR was performed on a Bio-Rad CFX Touch using SYBR Green PCR master mix (Roche) and primers for STING1 (Fwd; ATATCTGCGGCTGATCCTGC, Rev; TTGTAAGTTCGAATCCGGGC) and GAPDH (Fwd; CAGCCTCAAGATCATCAGCA, Rev; TGTGGTCATGAGTCCTTCCA). Samples were heated at 95°C for 5 mins before 44 cycles of 10 seconds at 95°C, 10 seconds at 60°C, 20 seconds at 72°C. Expression was normalised to *GAPDH*.

### RNA-seq library preparation and analysis

Total RNA was extracted using the RNeasy mini kit (Qiagen). Library preparation was carried out by the Edinburgh Clinical Research Facility from 500 ng of each RNA sample using the NEBNext Ultra II Directional RNA library kit with PolyA enrichment module (New England Biolabs). Libraries were assessed for quality and fragment size using the Agilent Bioanalyser. Sequencing was performed on the NextSeq 2000 platform (Illumina) using NextSeq 1000/2000 P2 Reagents.

FastQC was used to obtain basic quality control metrics from sequencing data and assess the quality of reads. For each sample, raw Fastq files were merged and aligned to the genome (hg19) using HISAT2 (Kim et al., 2019). Alignment statistics were calculated using GATK (Van der Auwera and O’Connor, 2020). Reads were assigned to genomic features using the featureCounts tool from the subread package (Liao et al., 2014).

Differential expression analysis between each set of conditions was carried out using DeSeq2 (Love et al., 2014). Contrasts were carried out between samples, where the sample specifies both the cell line and siRNA treatment. Gene ontology analysis was carried out using clusterProfiler (Wu et al., 2021). Volcano plots were rendered using ggplot2 (Wickham, 2016). A list of intronless genes was obtained from the hg19 GTF file available from UCSC (Nassar et al., 2023) by sorting for genes with a single exon. A list of histone genes was obtained from HGNC (Braschi et al., 2019).

For the analysis of transposable element expression, raw reads were aligned to the human genome assembly hg38 using HISAT2 (Kim et al., 2019). Alignment files were processed using the TEcounts tool from the TEtranscripts pipeline (Jin et al., 2015). Resulting transposable element and gene raw counts were then subjected to the variance stabilizing transformation in DESeq2 (Love et al., 2014) and analysed for differential expression with default settings.

### Statistics

Statistical analysis was performed using R and the specific statistical tests used are described in the relevant text and Figure legends. p-value significance is denoted as follows: * < 0.05, **< 0.01, *** < 0.001.

## Data availability

RNA-seq and ATAC-seq data generated in this study are being deposited at NCBI GEO.

## Author contributions

CRediT author statement:

W.A.B. – Conceptualisation, Funding acquisition, Project administration, Resources, Supervision, Writing – Original draft and revisions

B.M.B. – Conceptualisation, carrying out of experiments, data and statistical analyses, figure preparation, Writing – Original draft and revisions.

S.B. – Assisted with imaging of CCFs.

Y.K – Optimized ATAC-seq on IMR90 cells, prepared ATAC-seq libraries and analysed ATAC-seq data.

C.B. – Performed TPR KD for ATAC-seq, Writing – Revisions.

T.C. – Performed transposable element analysis.

J.C.A. – Conceptualisation, Supervision, Writing – Original draft and revisions.

A.Q. – Assisted with cell culture experiments.

## Acknowledgements

We thank the Edinburgh Clinical Research Facility for RNA-seq library preparation and for the sequencing of RNA-seq and ATAC-seq libraries. We thank the IGC Advanced Imaging facility for their help in fluorescence imaging and image analysis.

---

**Table S1.** Summary of day 8 ATAC-seq changes in peak accessibility. Number of peaks with significant changes generated from comparisons between samples using the limma package with an adjusted p-value cut-off of 0.05. Peaks significantly upregulated in RAS siCTRL compared to STOP siCTRL (SEN^+^) were further divided into TPR-dependent (SEN^+^TPR^+^) and TPR-independent (SEN^+^TPR^-^) as shown.

| Comparison | No. Peaks with significant changes |  |  |
| --- | --- | --- | --- |
| STOP siCTRL vs RAS siCTRL | 15732 | Down in RAS siCTRL: 8906 |  |
|  |  | Up in RAS siCTRL: 6826 | Down in RAS siTPR (SEN<sup>+</sup>TPR<sup>+</sup>): 1187 |
|  |  |  | No change in RAS siTPR (SEN<sup>+</sup>TPR<sup>-</sup>): 5639 |
|  |  |  | Up in RAS siTPR: 0 |
| STOP siCTRL vs STOP siTPR | 0 |  |  |
| RAS siCTRL vs RAS siTPR | 6185 | Down in RAS siTPR: 3972 |  |
|  |  | Up in RAS siTPR: 2213 |  |

**Table S2.** Summary of day 3 ATAC-seq changes in peak accessibility. Number of peaks with significant changes generated from comparisons between samples using the limma package with an adjusted p-value cut-off of 0.05.

| Comparison | No. Peaks with significant changes |  |
| --- | --- | --- |
| STOP siCTRL vs RAS siCTRL | 94394 | Down in RAS siCTRL: 58200 |
|  |  | Up in RAS siCTRL: 36194 |
| STOP siCTRL vs STOP siTPR | 0 |  |
| RAS siCTRL vs RAS siTPR | 0 |  |

**Table S3.**
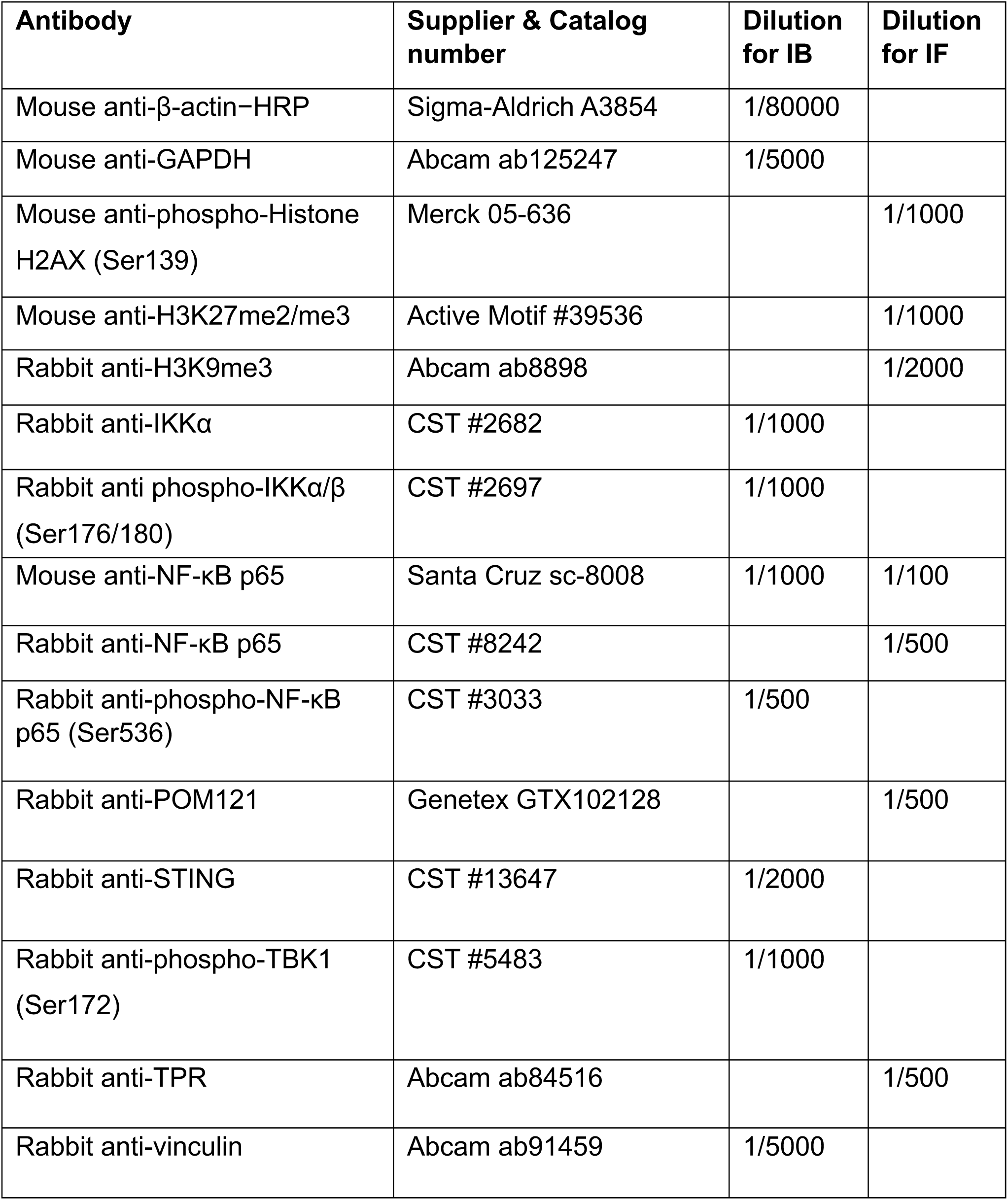
List of antibodies. Antibodies used in immunoblotting (IB) and immunofluorescence (IF) experiments with their corresponding dilutions.

| **Antibody** | **Supplier & Catalog number** | **Dilution for IB** | **Dilution for IF** |
| --- | --- | --- | --- |
| Mouse anti-β-actin-HRP | Sigma-Aldrich A3854 | 1/80000 | |
| Mouse anti-GAPDH | Abcam ab125247 | 1/5000 |  |
| Mouse anti-phospho-Histone H2AX (Ser139) | Merck 05-636 |  | 1/1000 |
| Mouse anti-H3K27me2/me3 | Active Motif #39536 |  | 1/1000 |
| Rabbit anti-H3K9me3 | Abcam ab8898 |  | 1/2000 |
| Rabbit anti-IKKα | CST #2682 | 1/1000 | |
| Rabbit anti phospho-IKKα/$\beta$ (Ser176/180) | CST #2697 | 1/1000 | |
| Mouse anti-NF-κB p65 | Santa Cruz sc-8008 | 1/1000 | 1/100 |
| Rabbit anti-NF-κB p65 | CST #8242 | | 1/500 |
| Rabbit anti-phospho-NF-κB p65 (Ser536) | CST #3033 | 1/500 | |
| Rabbit anti-POM121 | Genetex GTX102128 |  | 1/500 |
| Rabbit anti-STING | CST #13647 | 1/2000 |  |
| Rabbit anti-phospho-TBK1 (Ser172) | CST #5483 | 1/1000 |  |
| Rabbit anti-TPR | Abcam ab84516 |  | 1/500 |
| Rabbit anti-vinculin | Abcam ab91459 | 1/5000 |  |

